# Testing the Effect of the Toba Volcanic Eruption on Population Sizes in Worldwide Mammal Species

**DOI:** 10.1101/2020.04.06.028050

**Authors:** Nicole S. Torosin, Jennifer A. Raff, M. Geoffrey Hayes

## Abstract

The volcanic eruption of Toba in northern Sumatra at 71 kyBP (±5 kyBP) emitted sulfur gas and deposited thick layers of dust throughout the surrounding region. It is thought to have had a significant and dramatic cooling impact on the paleoclimate worldwide. Ambrose [1] conjectured this to be the cause of the contemporaneous (50-100 kyBP) population bottleneck observed in humans. We hypothesize that a volcanic winter of sufficient magnitude to cause a population bottleneck in humans would similarly affect other mammals. To test this hypothesis, we estimated pairwise mismatch distributions using mtDNA control region sequences of 28 mammal species archived on NCBI to assess whether each species underwent a population bottleneck. For any species fitting the sudden expansion model, we estimated the timing of the bottleneck and compared it to the date range of the Toba eruption. Only 3 of the 28 species show evidence of rapid population expansion overlapping in time with the Toba eruption. Therefore, the hypothesis that the volcanic winter triggered by the Toba eruption caused a significant bottleneck impacting mammal species worldwide is not supported by mitochondrial evidence. Our results question the hypothesis that the Toba eruption contributed to the bottleneck observed in humans at this time.

## 1 Introduction

A population bottleneck, defined as a drastic reduction in population size, reduces the genetic variation of a species resulting in founder effects [24]. Bottlenecks can be caused by events such as migration of small groups away from their larger population, landscape movement, or natural disasters on a local or global scale. Significant population decline can diminish genetic diversity and allow slightly deleterious alleles to increase in frequency [24].

The two largest bottlenecks in human prehistory occurred first after small populations dispersed out of Africa, and second, after a small group crossed the Bering land bridge into the Americas [18]. Evidence of these bottlenecks is reflected by the reduced genetic diversity and linkage disequilibrium that can be observed in the analysis of whole genomes [18]. The “Out of Africa” bottleneck was inferred by the observation of contrasting patterns of genetic diversity within and outside of Africa. The greatest genetic diversity occurs in African populations, where modern Homo sapiens originated [5, 18]. Heterozygosity of populations decreases with increased distance from Africa [2]. As small groups of *Homo sapiens* migrated out of Africa, each represented a small genetic sampling of the African population diversity, thereby creating founder effects.

A variation of the “Out of Africa” model is the “Weak Garden of Eden” hypothesis which posits that modern humans, after migrating to separate regions from Africa, underwent independent population bottlenecks and subsequent expansions [10]. Bottlenecks, and subsequent population expansion, can be detected through pairwise frequency histograms which graphically represents the number of differences in any two sequences within a set of DNA sequences [21]. Cann and colleagues [5] first created a pairwise distribution histogram by using restriction fragment length polymorphism (RFLP) patterns of 147 mtDNA extracts from various human populations: African American, aboriginal South African, Asian, European, aboriginal Australian, and aboriginal New Guinean. The RFLP haplotypes showed a bell-shaped unimodal distribution. Through simulation studies, Rogers and Harpending [21] showed that a bell-shaped unimodal distribution fit a model of sudden expansion. Using the pairwise distribution histograms, they estimated that all populations worldwide underwent sudden expansion at approximately 60-120 kyBP [21]. Later, Ambrose [1] notes that the release from the individual population bottlenecks worldwide is contemporaneous with the approximated time of climate warming post-volcanic winter, and with the transition from archaic to modern human technology found in the archaeological record [1]. As a result, he postulated that this observed global human bottleneck could have been caused by the super-eruption of the Toba volcano. The volcanic eruption of Toba in northern Sumatra at 66-76 kyBP emitted sulfur gas and deposited thick layers of dust throughout the region [6]. Further, it is thought to have had a dramatic cooling effect on the paleoclimate worldwide. Greenlandic ice core samples spanning this time period show high levels of sulfate and calcium, low levels of oxygen-18, and low solid ice electrical conductivity suggesting a massive volcanic eruption leading to a six-year volcanic winter, characterized by a severe and unprecedented cooling due to the obstruction of the sun by sulfuric acid droplets [33]. The volcanic winter was followed by the onset of a glaciation period which may have been independent of the Toba eruption but the high deposition of sulfuric acid just after the eruption indicates that the aerosols exacerbated the cooling process [33, 20]. The combination of events would cause drastic cooling precipitating the decimation of tropical plants and severe reduction of deciduous and temperate forests [11]. Drop in temperature and loss of food sources would severely impact fauna worldwide, possibly leading to substantial population reductions. According to this hypothesis, populations in equatorial Africa would have been less affected by the global cooling, allowing African populations to maintain higher population sizes and greater genetic diversity [1].

Other geological studies have attenuated Ambrose’s [1] hypothesis. For example, primate species living in the rainforest habitat surrounding Toba do not appear to have experienced bottlenecks as shown by fossil data [15] and the sediment of Lake Malawi, 7,000 km west of Toba in East Africa, shows no major change in sediment composition or temperature due to the Toba eruption [14]. To better explain the timeline of human evolution and evaluate whether this paleoclimatic event had a disparate effect on the genetic than the geological record, we decided to genetically test Ambrose’s [1] hypothesis that the Toba volcanic winter is coincident with the timing of the modern human bottlenecks, population size reduction, and subsequent expansion of anatomically modern humans. We postulate that a global volcanic winter would have affected other mammals worldwide in addition to *H. sapiens*.

To test our hypothesis, we collated publicly available mtDNA hypervariable segment (HVS) sequences from 28 mammal species spanning the globe, created pairwise mismatch distribution histograms, and estimated the timing of any bottlenecks found. The HVS is a non-coding region allowing synonymous mutations to accumulate [25]. This feature makes the HVS region ideal for detecting sudden expansion. We understand the limitations of using modern mtDNA samples to infer population history and that it would be beneficial to combine our analysis with microsatellites or ancient DNA, however, recent studies continue to use mtDNA to evaluate demographic history [17, 4, 28, 32].

## 2 Results

This study evaluated the mtDNA control region of 28 mammalian species for evidence of population bottlenecks followed by sudden expansion post-super-eruption of the Toba volcano in Sumatra (Figure 1). By creating pairwise mismatch distribution histograms, we were able to distinguish which histograms fit Rogers and Harpending’s [21] model of sudden expansion. We evaluated pairwise distribution histograms for waves characteristic of sudden expansion. Figure 2 highlights plots where the mtDNA data converges to the pattern predicted by Arlequin to be sudden expansion. Thirteen mammalian species including *Homo sapiens* exhibited a demographic bottleneck in their evolutionary history.

**Figure 1:**
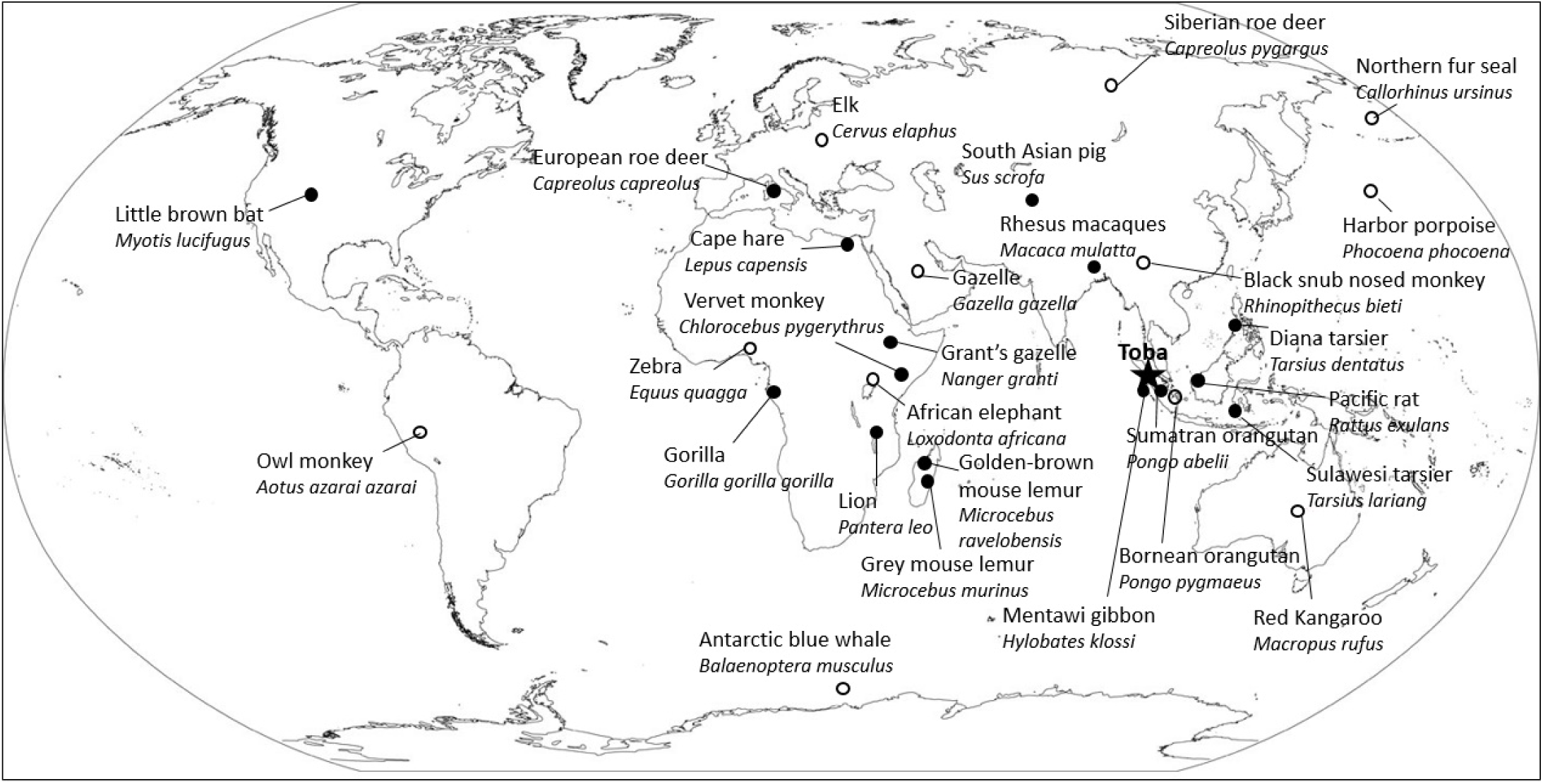
Map pinpointing geographical location of all species analyzed. Closed circle: no bottleneck visible in the pairwise mismatch distribution histogram. Open circle: bottleneck evident in the pairwise mismatch distribution histogram.

**Figure 2:**
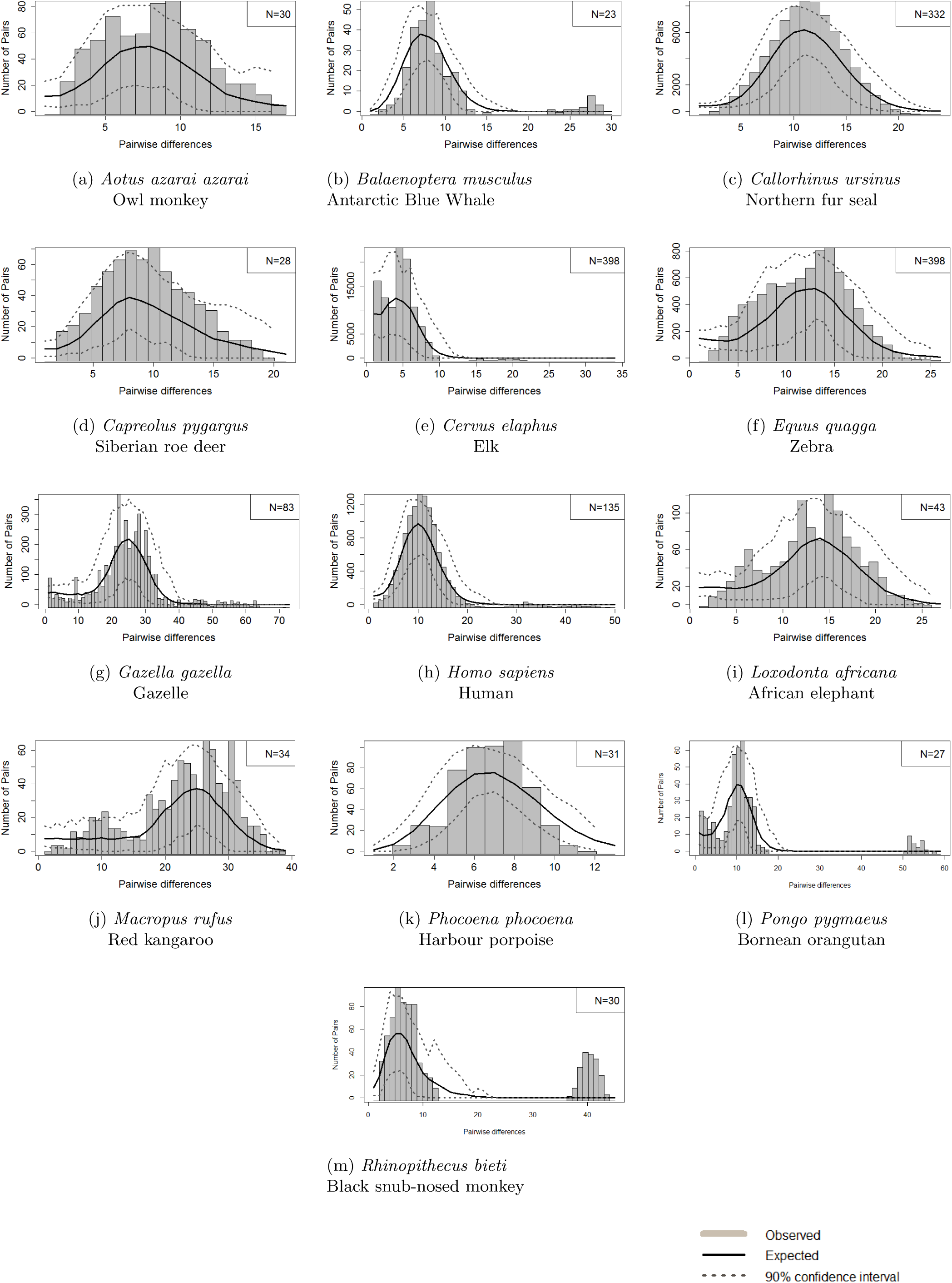
Pairwise mismatch distribution histograms of species showing bottlenecks in evolution. Gray bars: observed pairwise mismatch distribution, Black lines: Arlequin simulated distribution of sudden expansion, Dotted line: 90% confidence interval.

**Figure 3:**
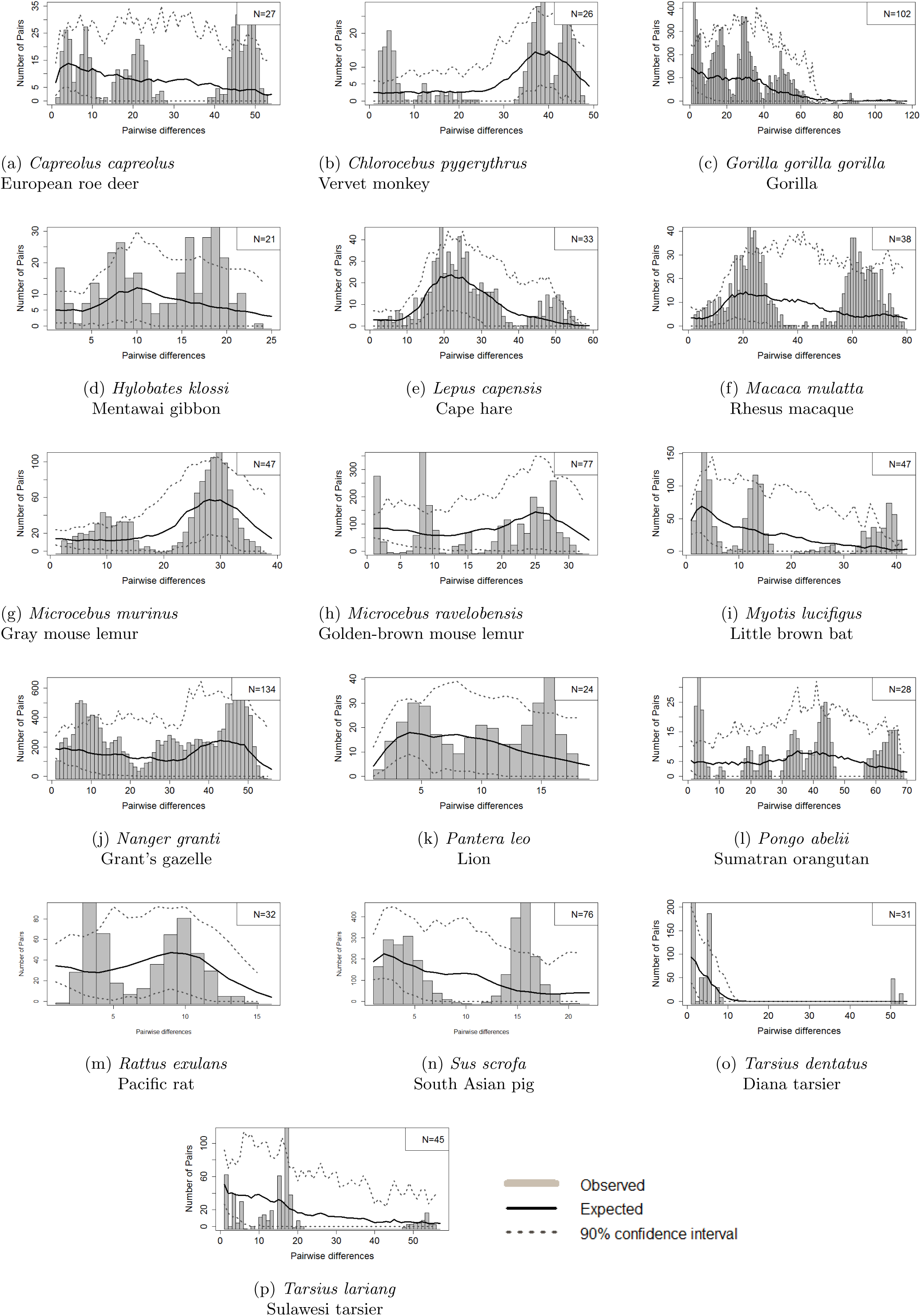
Pairwise mismatch distribution histograms of species showing equilibrium in evolution. Gray bars: observed pairwise mismatch distribution, Black lines: Arlequin simulated distribution of equilibrium, Dotted line: 90% confidence interval.

Larger p-values and raggedness index p-values indicate that the form of the observed pairwise mismatch distribution histogram fits the sudden expansion model. Irregularity in the pairwise mismatch distribution due to sites prone to polymorphisms or bimodal distributions prevented the use of the p-values to support a bottleneck. For example, the p-value and raggedness index p-value for *Gazella gazella* and *Balaenoptera musculus* are both zero, or close to zero, because the observed pairwise mismatch distribution is irregular and does not fit the expected mismatch distribution. However, the shape of the histogram, seen more clearly in the curves of the 90% confidence intervals, follows the unimodal trend revealing a bottleneck.

We also assessed the mtDNA alignments for Tajima’s D and Fu’s Fs neutrality statistics (Table 1 and Table 2). The Fu’s Fs statistics for all 13 mammals displaying bottlenecks are negative, indicating excess diversity of alleles, as expected during a sudden population expansion. The Tajima’s D statistics are all negative, indicating an excess of low-frequency polymorphisms. Excess of low frequency polymorphisms represents regrowth of segregating sites and heterozygosity after genetic depletion, as expected during a bottleneck followed by population expansion.

**Table 1:** Data for species fitting the sudden expansion model.

| Species | Common name | p-value | Raggedness index | p-value | $\theta_0$ | $\theta_1$ | Fu's $F_s$ | p-value | Tajima's $D$ | p-value |
| --- | --- | --- | --- | --- | --- | --- | --- | --- | --- | --- |
| *B. musculus* | Antarctic blue whale | 0.02 | 0.264 | 0 | 0 | 99999 | -26.96 | 0.007 | -2.95 | 0.989 |
| *C. pygargus* | Siberian roe deer | 0.73 | 0.048 | 0.61 | 0 | 99999 | -26.58 | 0.018 | -2.58 | 0.998 |
| *C. ursinus* | Northern fur seal | 0.05 | 0.011 | 0.11 | 0.046 | 124.47 | -24.31 | 0.089 | -1.34 | 0.871 |
| *R. bieti* | Black snub nosed monkey | 0.06 | 0.053 | 0.5 | 0 | 314.4 | -25.31 | 0.02 | -2.01 | 1 |
| *L. capensis* | Cape hare | 0.9 | 0.005 | 0.63 | 10.32 | 213.95 | -15.81 | 0.012 | -0.53 | 0.998 |
| *M. murinus* | Gray mouse lemur | 0.02 | 0.028 | 0 | 0.007 | 49.34 | -24.3 | 0.054 | -0.72 | 1 |
| *L. africana* | African elephant | 0.12 | 0.037 | 0.07 | 0 | 49.45 | -25.54 | 0.079 | -1.86 | 1 |
| *E. quagga* | Zebra | 0.05 | 0.037 | 0.05 | 0 | 28.96 | -25.49 | 0.163 | -1.72 | 1 |
| *G. gazella* | Gazelle | 0 | 0.04 | 0 | 0.004 | 71.8 | -24.15 | 0.196 | -0.18 | 1 |
| *H. lar* | Lar gibbon | 0.25 | 0.13 | 0.3 | 0.004 | 99999 | -5.38 | 0.432 | -3.07 | 0.995 |
| *M. rufus* | Red kangaroo | 0.49 | 0.008 | 0.76 | 0 | 52.27 | -24.57 | 0.001 | -1.59 | 0.998 |
| *P. pygmaeus* | Bornean orangutan | 0.04 | 0.102 | 0.02 | 0.002 | 21.33 | -25.41 | 0.022 | -2.44 | 0.989 |
| *H. sapiens* | Human | 0.61 | 0.005 | 0.63 | 0.08 | 97.66 | -24.41 | 0.067 | -1.78 | 0.653 |

**Table 2:**
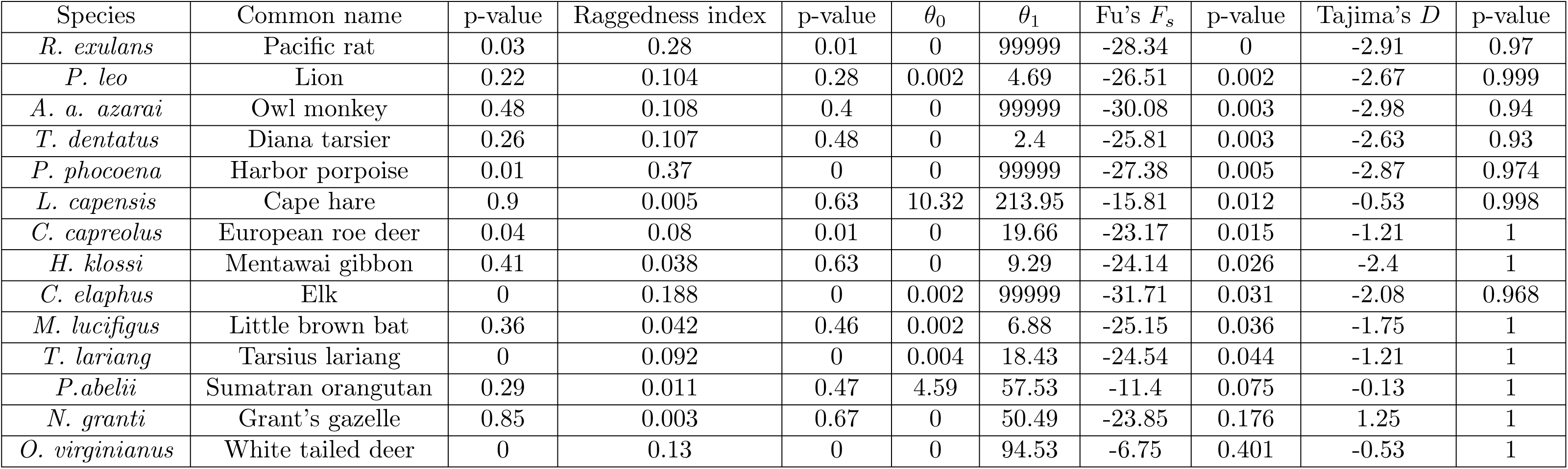
Data for species in equilibrium.

Using the tau values we were able to calculate the time since sudden expansion. Table 3 contains species showing sudden expansion due to bottleneck and outlines the information necessary to calculate the time since expansion of each species’ bottleneck: tau values, nucleotide substitution rate, generation time, and base pair length of mtDNA. The range of time since expansion estimated for two species in addition to *Homo sapiens* corresponded with the eruption of Toba: *Capreolus pygargus*, with a median time since sudden expansion estimated at 73,071 years ago, and *Rhinopithecus bieti* with an upper bound of time since sudden expansion estimated at 63,913 years ago.

**Table 3:**
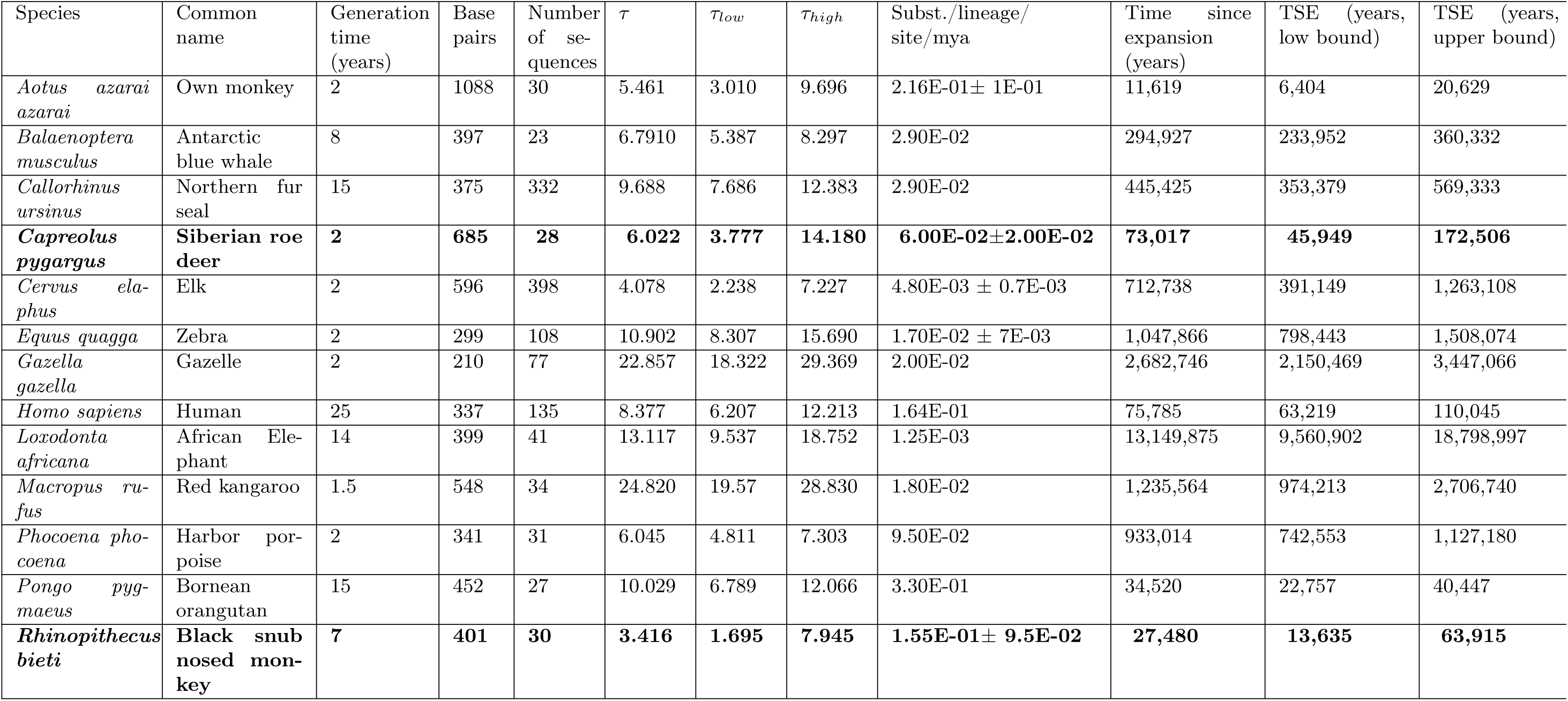
Species fitting the sudden expansion model based on pairwise mismatch distribution histograms. Bolded species are those with time since expansion overlapping in time with the Toba eruption.

## 3 Discussion

Humans worldwide underwent a population bottleneck 50,000-100,000 years ago [21, 5]. Ambrose [1] hypothesized that the bottleneck was a result of the volcanic winter caused by the super-eruption of the Toba volcano 71 kyBP (±5 kyBP) [6, 33]. In order to test Ambrose’s theory, we supposed that if humans underwent a drastic population reduction as a result of the Toba eruption, other mammalian species, especially those near Sumatra and outside the African area of refugia, would have as well. We compiled sets of control region mammalian mtDNA and subjected them to bottleneck analysis. Worldwide, bottlenecks were seen in 12 out of the 28 mammalian species we tested in addition to humans.

The results do not support the Toba eruption as the root cause of the bottleneck observed in humans. While almost half of the species exemplified a bottleneck in their evolution, the timing of the bottleneck of only two species examined, *Capreolus pygargus* and *Rhinopithecus bieti*, could correspond to the mega-eruption of the Toba volcano.

Primates *Pan abelii, Hylobates klossi, Tarsius dentatus*, and *Tarsius lariang*, native to the rainforest habitat surrounding Toba, which would have been highly depleted by a global winter, do not appear to have been experienced bottlenecks at all. This finding has been independently confirmed with fossil data [15]. Considering that species predicted to be highly affected by the global winter due to their reliance on tropical vegetation were not affected, the global winter caused by the Toba eruption was potentially not as severe as previously thought.

A study on the sediment of Lake Malawi, 7,000 km west of Toba in East Africa, showed no major change in sediment composition or temperature due to the Toba eruption [14]. This geological evidence, unearthed more than 10 years after Ambrose’s paper, reduces the strength of his conclusions. Compounded with the lack of bottleneck and extinction evidence [15] in species surrounding Toba, our genetic research is in agreement with geological evidence suggesting that the effect of Toba’s eruption was not as great as hypothesized by Ambrose [1]. Another possible explanation for the lack of genetic evidence of major bottlenecks coincident with the Toba eruption is that animal populations often recover quickly from catastrophic events [30, 7]. For example, *Capreolus pygargus* showed a bottleneck contemporaneous with the Toba eruption, but it is a cold-adapted species that species can rely on alternate food sources when necessary [12].

Another explanation of the lack of species showing evidence for a bottleneck is that the harsh effects of the eruption severely reduced the population sizes of the species within its vicinity, but post-disaster remigration into the area and intermixing with other populations obscured evidence of the bottleneck in the mtDNA record [26]. An ideal method to determine whether intermixing obscured evidence of a bottleneck in the pairwise mismatch distribution histograms is an analysis of phylogenetic networks for each species. This analysis would use haplotypes, specific combinations of alleles inherited together, present in the individual mtDNA sequences to determine common ancestry (monophyletic) or separate ancestry (paraphyletic). Paraphyletic phylogeny would imply that another population of the same or related species migrated into the Toba region and intermixed with the remaining local population, consolidating into one species. Pairwise mismatch distribution histograms for two species fitting the sudden expansion model were bimodal, representing two population bottlenecks in their evolutionary history. Based on observation, the second mode in a bimodal distribution, representing the older bottleneck, causes tau value calculation of the younger bottleneck to be marginally higher than the tau values of unimodal pairwise distributions. While mammals with bimodal pairwise mismatch distributions may have had a slightly higher tau value, we do not believe this would drastically alter the estimations of time since expansion. Presently, population expansion simulations testing the effects of two bottlenecks have not been researched and published to permit a full understanding of how to determine the distinct tau value of each bottleneck visible in the histograms.

Lastly, using mtDNA solely cannot always detect bottlenecks or their timing with certainty [17]. Further research will help test the conclusions based on mammalian mtDNA analysis in this study. These additional approaches can include pairwise mismatch distribution analysis of plant or chloroplast DNA, testing the mtDNA cytochrome B region or nuclear DNA of mammals used in this study, and expanding the study to include other taxa. If additional analyses find that the Toba eruption did not have a substantial effect on all flora and fauna, other explanations for the simultaneous bottleneck releases of human populations worldwide described by Cann and Rogers and Harpending [21, 5] should be tested.

## 4 Methods

We assembled mtDNA sequences from the non-coding control region (a.k.a. D-loop, hypervariable region or hypervariable segment) from 28 mammalian species spanning the world (Table 1, Table 2, Figure 1). The control region of mtDNA is an ideal resource for bottleneck identification through pairwise mismatch distribution calculation because the sequence is non-coding and non-recombining [29]. Non-coding regions allow putatively neutral mutations to accumulate and the bottleneck to be dated using the number of mutations and the known nucleotide substitution rate of the mtDNA of the species [25]. We searched the National Center for Biotechnology Information Search database (NCBI) for publications that sequenced a minimum of 25 samples for the species of interest. We selected datasets for species within one logarithmic order of human size to limit the variation of nucleotide substitution rate [16]. The global human weight average is 62 kg [31] which allowed for species between 22.8 - 169 kg to be used in our analysis. We compiled species-specific nucleotide substitution rates in terms of nucleotide substitutions per lineage per site per mya from the literature (Supplemental Table 1). We aligned mtDNA sequences using MAFFT v.7 [13] multiple sequence alignment program. Individual mtDNA sequences with >5% missing nucleotide calls were also removed [3]. GENBANK or ENA accession numbers for samples included are listed in Supplementary Table 1.

We input mtDNA alignments into Arlequin ver 3.5.1.2 [8] and selected “Estimate parameters of demographic expansion” under the “Mismatch Distribution” tab within “Arlequin Settings” with 100 bootstrap replicates to obtain observed and expected mismatch distributions and associated values including sum of squared deviations and p-value, raggedness index and p-value, and tau values, [21, 23]. Additionally, we used Arlequin ver 3.5.1.2 [8] and selected “Fu’s Fs” with 1000 simulations [9] and “Tajima’s D” under “Neutrality tests” [27] to obtain neutrality statistics for each alignment. We set deletion weight equal to 0, and transitions (Ts), transversions (Tv), indel weights equal to 1, and allowed “missing level per site” to 0.05 in the section “General Settings” (personal communication Laurent Excoffier and Alan Rogers). Parameter tau is a unit of mutational time represented by the modal number of pairwise differences and used to calculate the time since expansion using the formula, where T is the time since expansion, *τ* is tau, and *µ* is the nucleotide substitutions per lineage per site per generation [21]. Fu’s Fs measures the diversity of alleles [9], while Tajima’s D evaluates the presence of non-neutral evolution such as selection or rapid expansion [27]. Raggedness index indicates the similarity between the observed and simulated pairwise mismatch distribution models [10].

We created histograms of the observed and expected pairwise mismatch frequencies in R [19] by plotting the number of pairs with each number of differences in their sequences. Species with a genetic bottleneck are identified due by a smooth, unimodal wave in the mismatch distribution [21]. A species in equilibrium is described by a flat or scattered, patternless distribution [21]. Using this metric, we identified species with genetic bottlenecks in their history.

In order to calculate the time since post-bottleneck expansion for species with a bottleneck, we used the Schenekar and Weiss [22] mismatch calculator. Using the tau values, including error range based on 90% confidence interval for tau values and correcting for sequence length and generation time, we calculated time since expansion for the thirteen species showing evidence of sudden expansion based on their pairwise mismatch distribution histograms.

