## Supplemental Table 1 for "Testing the Effect of the Toba Volcanic Eruption on Population Sizes in Worldwide Mammal Species"

### Supplement

Table 1: References for mtDNA sequences and substitution rates. Note: Empty cells correspond to species showing equilibrium in evolution that did not require a substitution rate

| Species | Common name | mtDNA region | mtDNA Accession numbers | Reference | Substitution rate | Reference |
|---|---|---|---|---|---|---|
| *Aotus azarai azarai* | Owl monkey | Control region | JN161069-JN161098 | [2] | 2.16E-01± 1E-01 | [2] |
| *Balaenoptera musculus* | Antarctic blue whale | Control region | JN801048-JN801070 | [27] | 2.90E-02 | [12] |
| *Callorhinus ursinus* | Northern fur seals | Control region | EU791990-EU792321 | [6] | 2.90E-02 | [6] |
| *Capreolus capreolus* | European roe deer | D-loop | JX971589-JX971615 | [13] | | |
| *Capreolus pygargus* | Siberian roe deer | Control region | JX428900-JX428927 | [13] | 6.00E-02±2.00E-02 | [22] |
| *Cervus elaphus* | Elk | Control region | JX125702-JX126108 | [11] | 4.80E-03 ± 0.7E-03 | [7] |
| *Chlorocebus pygerythrus* | Vervet monkey | D-loop | KP231259-KP231284 | [9] | | |
| *Equus quagga* | Zebra | D-loop | EU650488-EU650599 | [16] | 1.70E-02 ± 7E-03 | [21] |
| *Gorilla gorilla gorilla* | Gorilla | Control region: HV1 | EU305296- EU305397 | [1] | | |
| *Homo sapiens* | Humans | D-loop: HV1 | M76235-M76368 | [29] | 1.64E-01 | [26] |
| *Hylobates agilis* | Agile gibbon | Control region:HV1 | EF363485-EF363509 | [32] | | |
| *Hylobates lar* | Lar gibbon | D-loop | AB720992-AB721002 | [17] | | |
| *Lepus capensis* | Cape hare | D-loop | HM233286-HM233263 | [14] | | |
| *Loxodonta africana* | African Elephant | Control region | AF106203-AF106245 | [20] | | |
| *Macaca mulatta* | Rhesus macaque | D-loop | JF746816- JF746845, JF834912-JF834921 | [34] | | |
| *Macropus rufus* | Red kangaroo | Control region | AJ225130- AJ225163 | [5] | 1.80E-02 | [5] |
| *Microcebus murinus* | Grey mouse lemurs | D-loop | JF796804-JF796850 | [25] | | |
| *Microcebus ravelobensis* | Golden-brown mouse lemur | D-loop | EF065308- EF065385 | [10] | | |
| *Myotis lucifugus* | Little brown bat | Control region: HV2 | EF471399-EF471445 | [28] | | |
| *Nanger granti* | Gazelles | Control region | EU029806-EU029940 | [16] | | |
| *Odocoileus virginianus* | White tailed deer | D-Loop | JN088006-JN088047 | [4] | | |
| *Pantera leo* | Lion | Control Region: HVR1 | DQ899900 - DQ899923 | [3] | | |
| *Phocoena phocoena* | Harbor porpoise | Control region | FJ214731-FJ214801 | [24] | | [30] |
| *Pongo abelii* | Sumatran orangutan | D-loop: HV1 | JQ962945-JQ962972 | [19] | | |
| *Pongo pygmaeus* | Bornean orangutan | D-loop | AJ391123, AJ391124-5, AJ391098, AJ391100-03, AJ391105-10, AJ391113-18, AJ391120-21, AJ391132-34, AJ391136.2-37.2 | [31] | 3.30E-01 | [31] |
| *Rattus exulans* | Pacific rat | D-loop | AY604202-AY604233 | [23] | | |
| *Rhinopithecus bieti* | Black snub nosed monkey | D-loop | DQ661651-DQ661680 | [15] | 1.55E-01± 9.5E-02 | [33] |
| *Sus scrofa* | South Asian pigs | Control region | JQ238237-JQ238602 | [8] | | |
| *Tarsius dentatus* | Diana tarsier | Control region: HV1 | FJ614477-FJ614489, FJ614493-FJ614494, FJ614497-FJ614500, FJ614502-FJ614509 | [18] | | |
| *Tarsius lariang* | Sulawesi tarsier | Control region: HV1 | FJ614403- FJ614444 | [18] | | |

[34] H. Z. Z. Y. L. S. Yu, W.H. Mitochondrial dna (mtdna) d-loop region complete sequence of macaca mulatta. *unpublished*, 2011.
